## Supplementary figures and images for "Gene expression profiling of malaria parasites reveals common virulence gene expression in adult first-time infected patients and severe cases"

### Figure S1

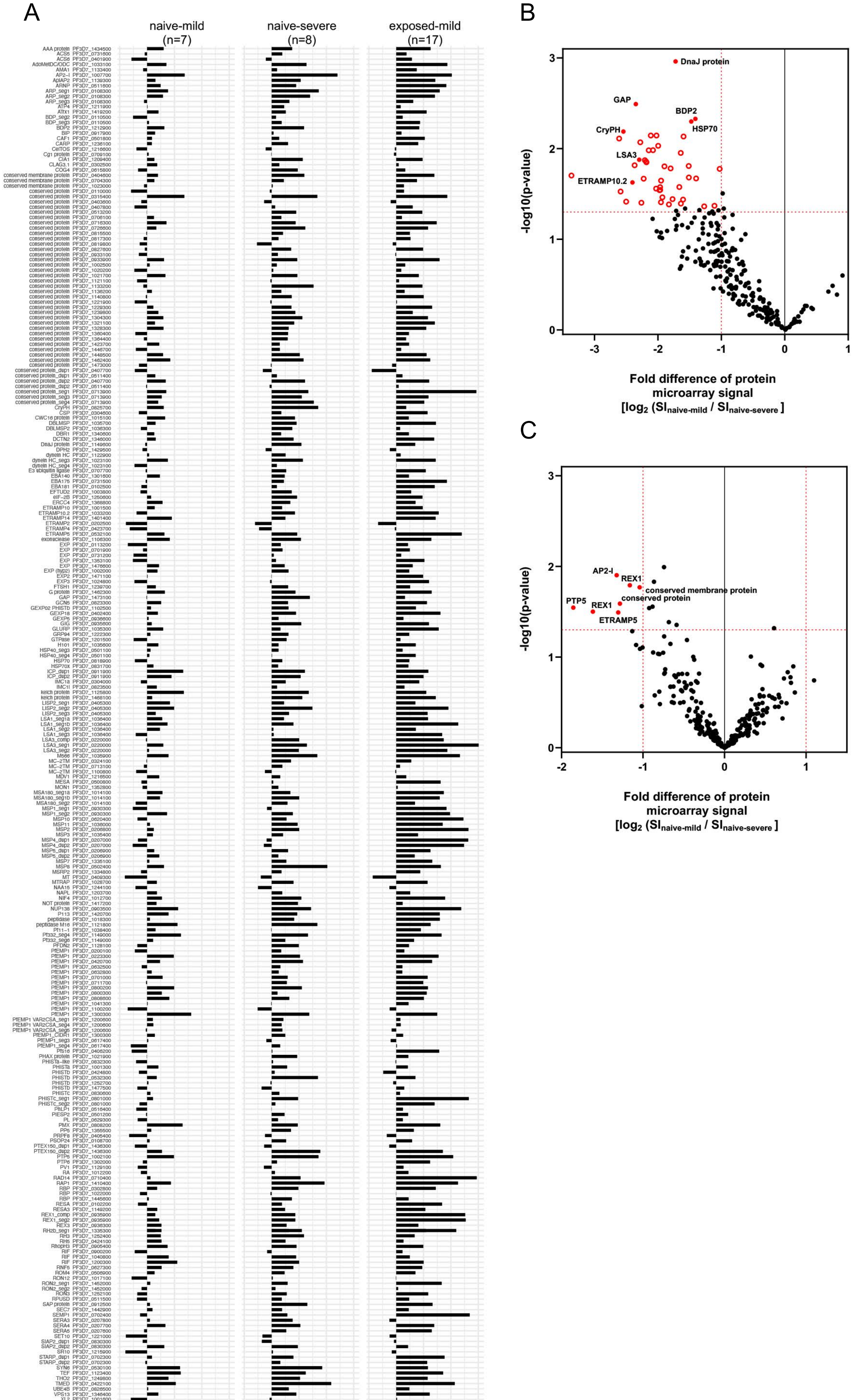

### Figure S2

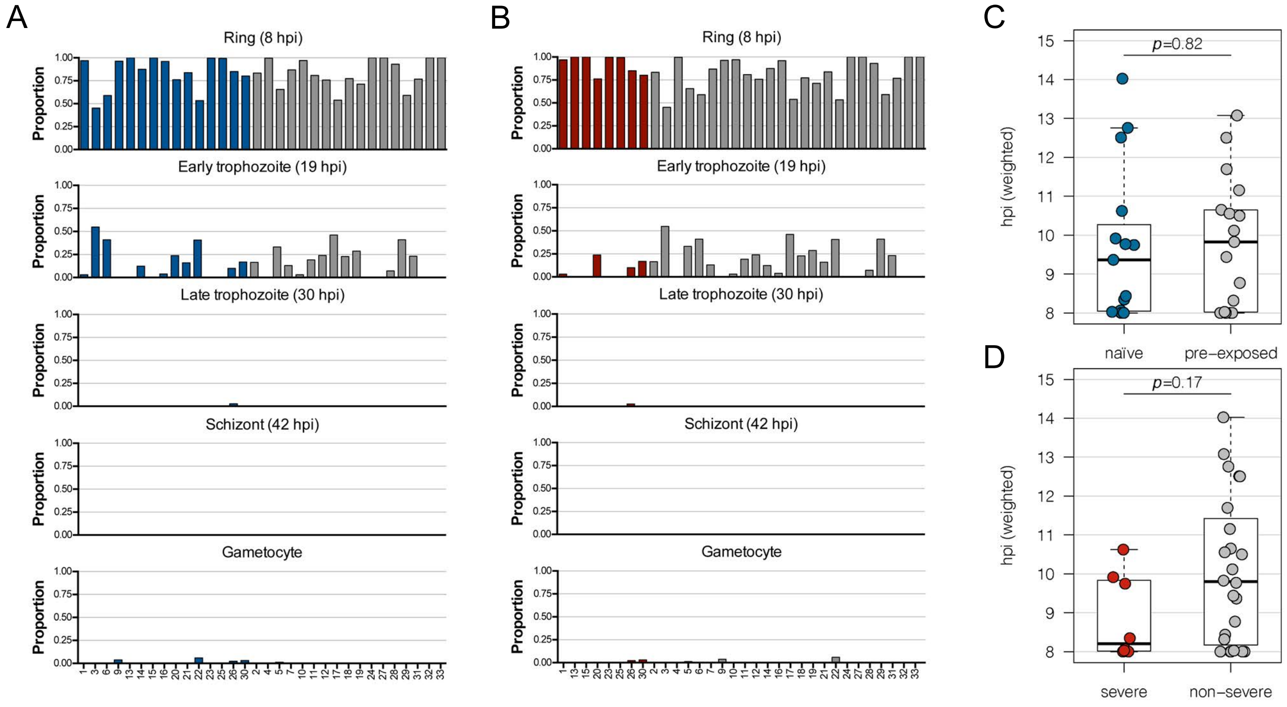

### Figure S3

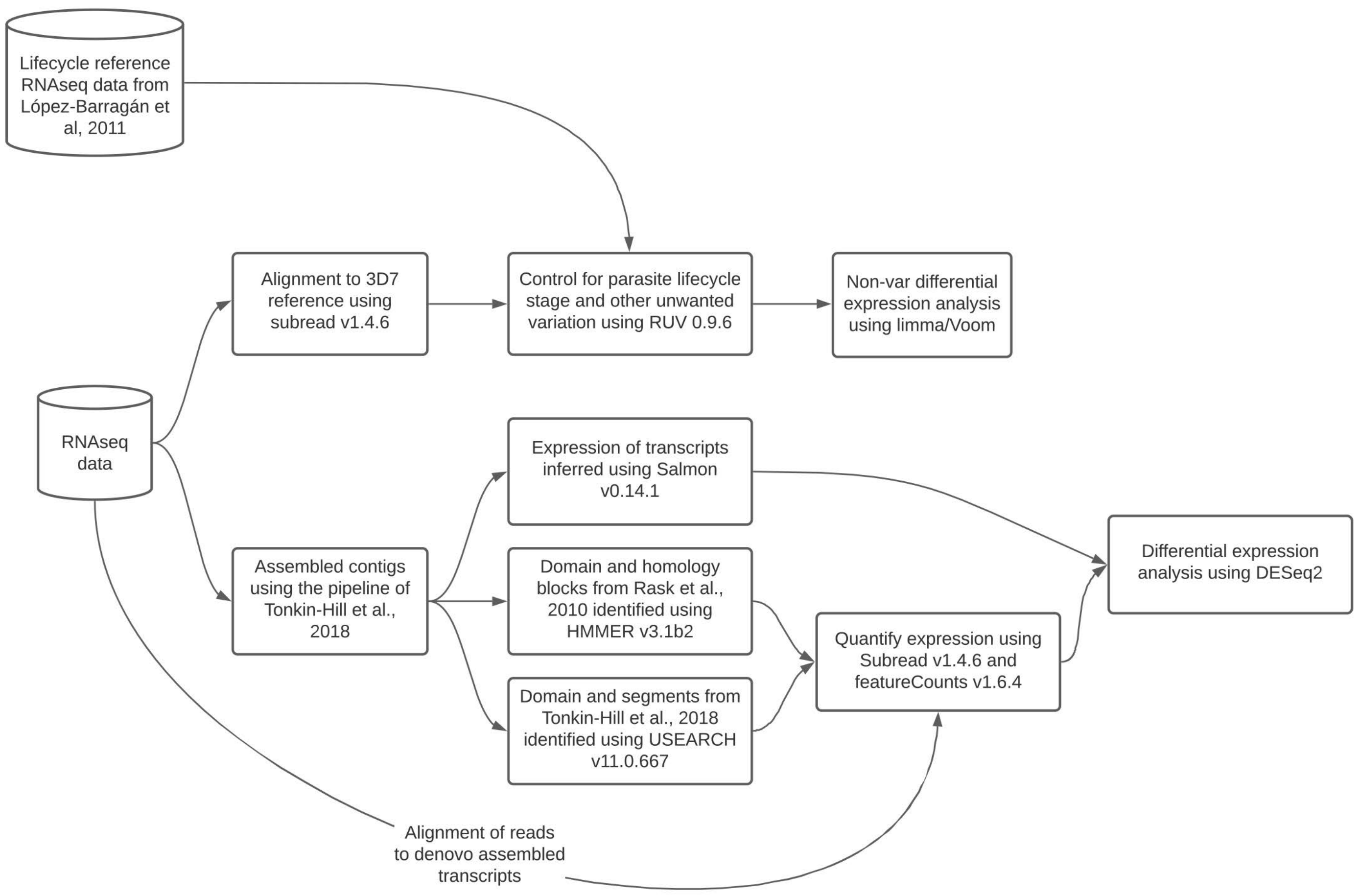

### Figure S5

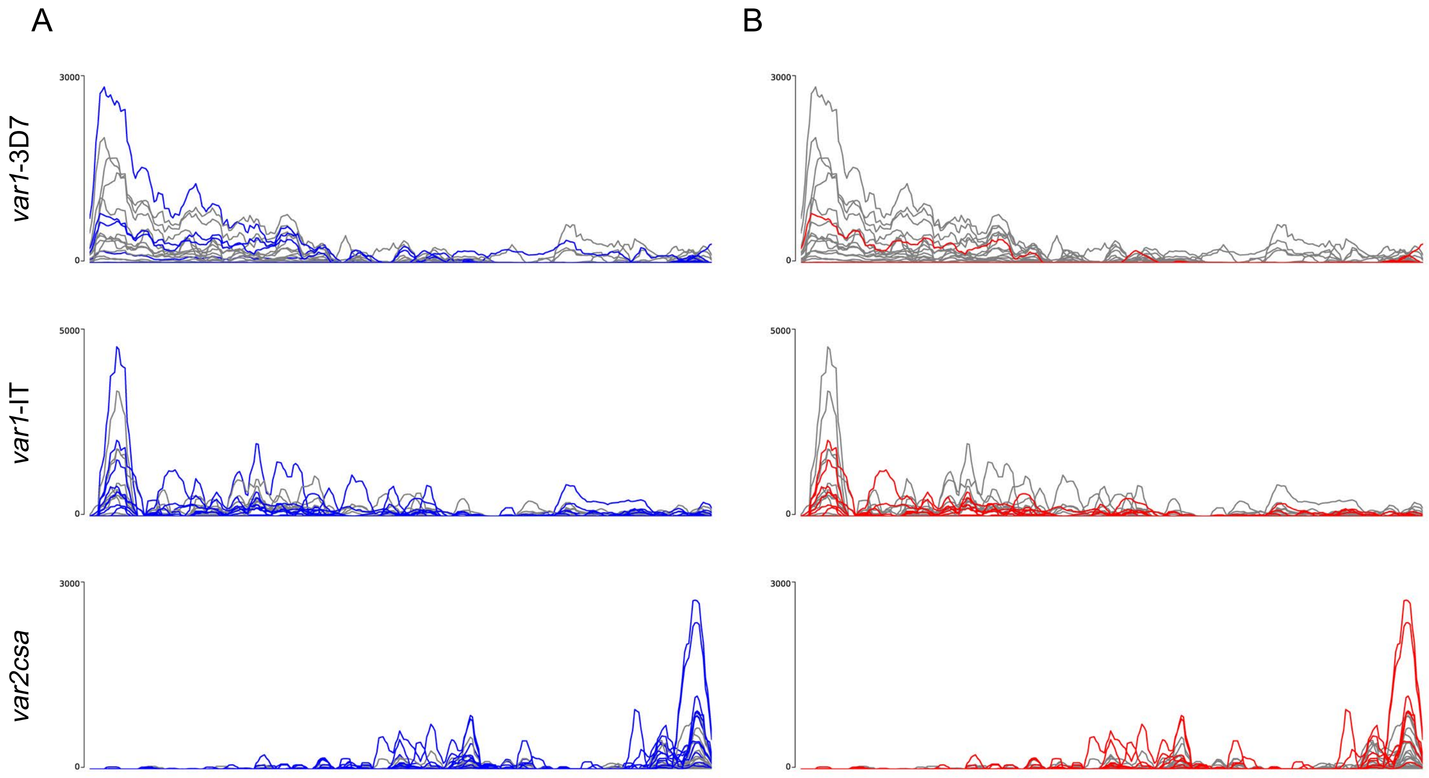

### Figure S6

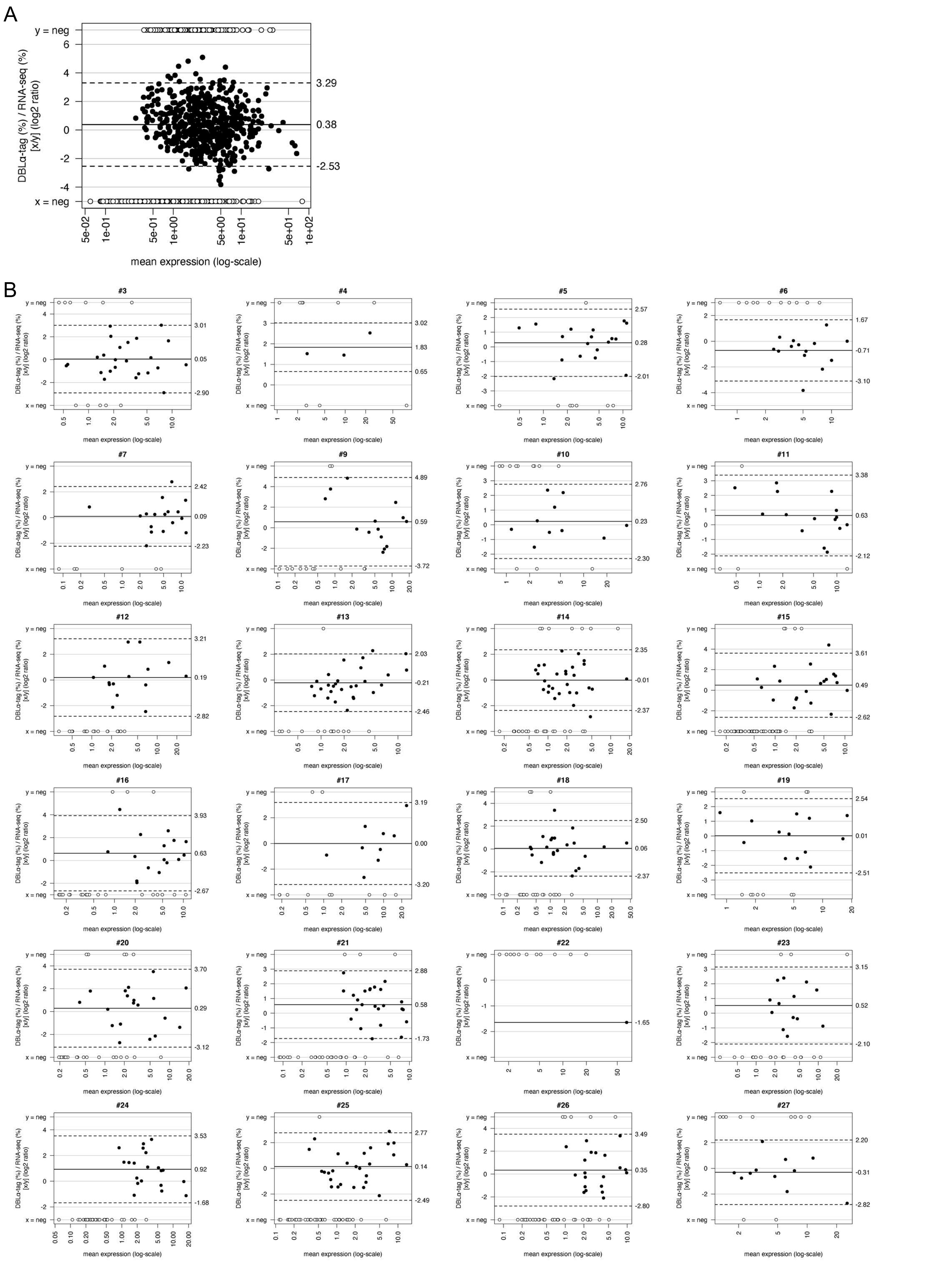

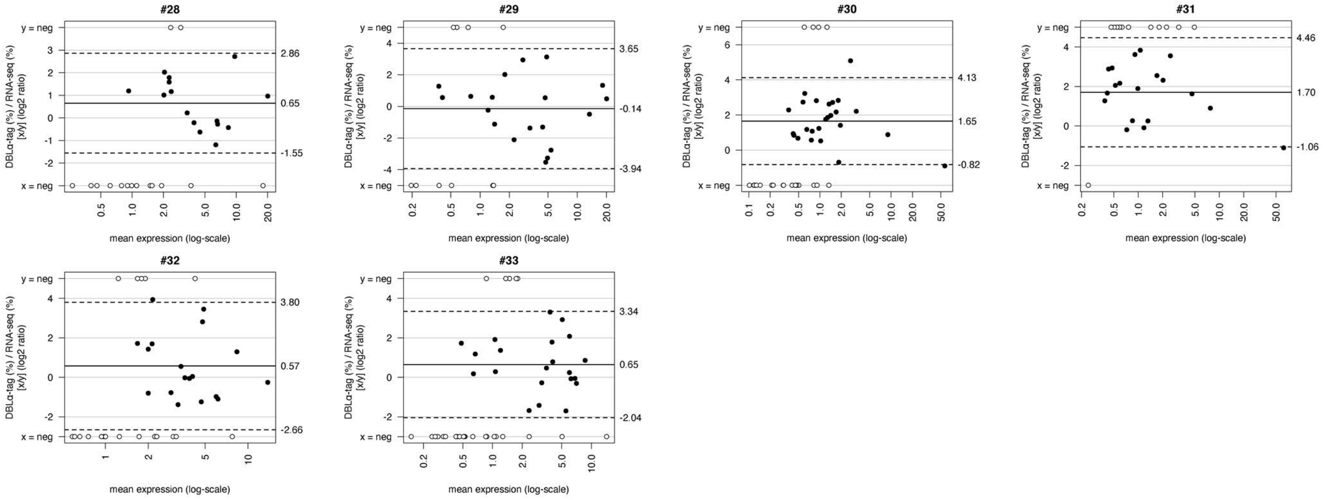

### Figure S7

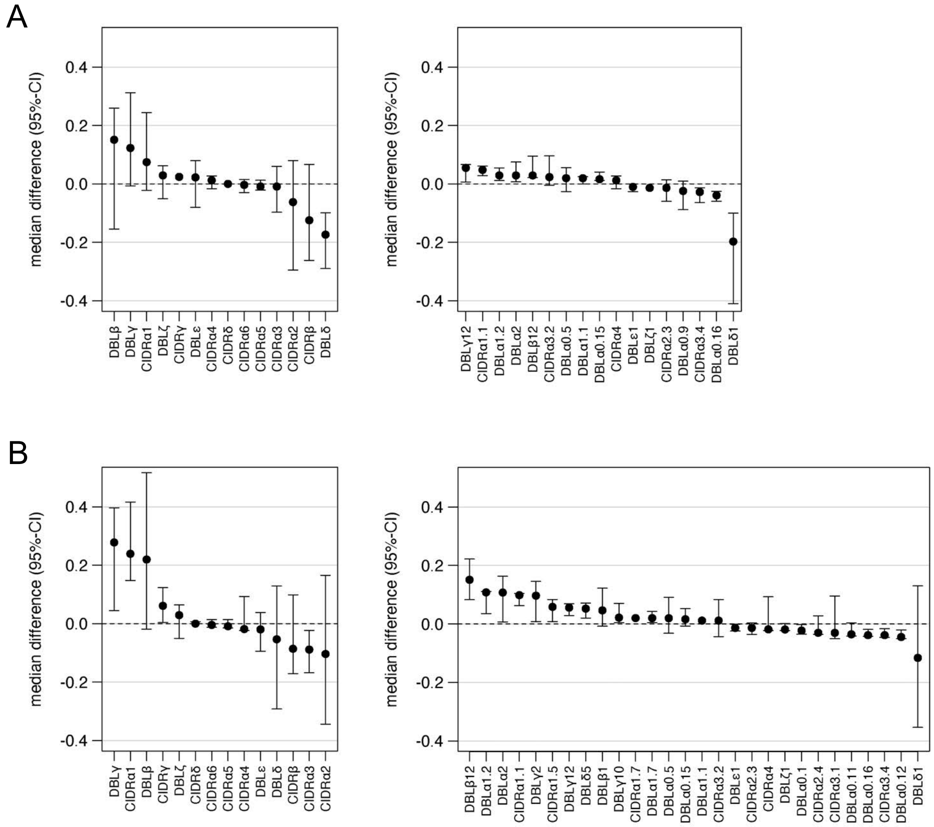

### Figure S8

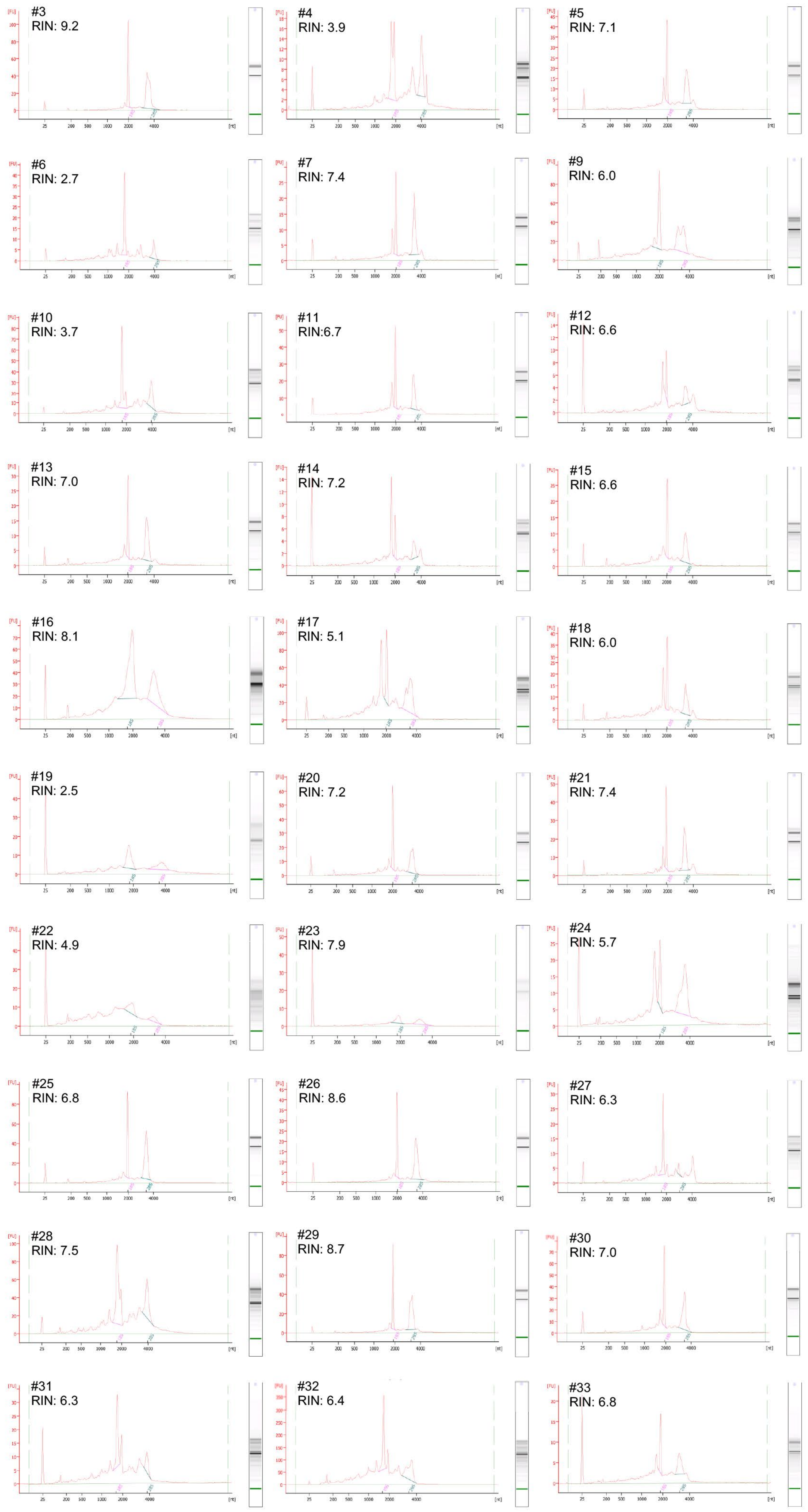
