## Supplementary material for "Gene expression profiling of malaria parasites reveals common virulence gene expression in adult first-time infected patients and severe cases": Figure S4

A

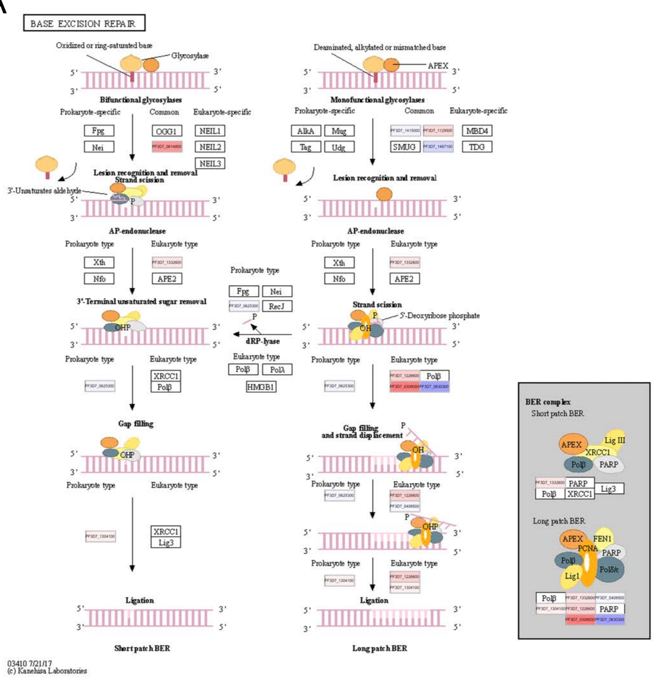


B

| GeneID | logFC | adj.P.Val | KEGG abreviation | [Product Description] |
|---|---|---|---|---|
| PF3D7_1361900 | 1,144 | **0,00723** | PCNA | proliferating cell nuclear antigen 1 |
| PF3D7_0614800 | 1,338 | **0,00634** | NTH | endonuclease III homologue, putative |
| PF3D7_0308000 | 1,617 | **0,00384** | Pold | DNA polymerase delta small subunit, putative |
| PF3D7_1415000 | 0,11 | 0,846 | UNG | uracil-DNA glycosylase |
| PF3D7_1129500 | 0,641 | **0,0121** | MUTY | A/G-specific adenine glycosylase, putative |
| PF3D7_1467100 | -0,201 | 0,359 | MPG | DNA-3-methyladenine glycosylase, putative |
| PF3D7_1332600 | 0,528 | **0,0336** | APE1/APEX | apurinic/apyrimidinic endonuclease Apn1, putative |
| PF3D7_0625300 | 0,128 | 0,731 | Dpol | DNA polymerase 1, putative |
| PF3D7_1411400 | 0,243 | 0,503 | Dpol | plastid replication-repair enzyme |
| PF3D7_1226600 | 0,724 | 0,0708 | PCNA | proliferating cell nuclear antigen 2 |
| PF3D7_1017000 | 0,961 | **0,0285** | Pold | DNA polymerase delta catalytic subunit |
| PF3D7_0630300 | -0,761 | 0,185 | Pole | DNA polymerase epsilon catalytic subunit A, putative |
| PF3D7_1234300 | 0,29 | 0,298 | Pole | DNA polymerase epsilon subunit B, putative |
| PF3D7_1304100 | 0,411 | 0,526 | Lig1 | DNA ligase I |
| PF3D7_0408500 | 0,152 | 0,723 | Fen1 | flap endonuclease 1 |
