## Supplementary material for "Gene expression profiling of malaria parasites reveals common virulence gene expression in adult first-time infected patients and severe cases": Table S2

**Table S2: Characteristics of each patient.**

| ID # | Parasitemia | Signs of organ failure | Classification |
| --- | --- | --- | --- |
| 1 | 1% | No* | Naïve, severe |
| 2 | 1% | No | Pre-exposed, non-severe |
| 3 | >5% | No | Naïve, non-severe |
| 4 | 1.5% | No | Pre-exposed, non-severe |
| 5 | 1.5% | No | Pre-exposed, non-severe |
| 6 | 0.5% | No | Naïve, non-severe |
| 7 | 2.5% | No | Pre-exposed, non-severe |
| 9 | 7% | No | Naïve, non-severe |
| 10 | 2% | No | Pre-exposed, non-severe |
| 11 | 3% | No | Pre-exposed, non-severe |
| 12 | 0.5% | No | Pre-exposed, non-severe |
| 13 | 35% | Yes | Naïve, severe |
| 14 | <1% | No | Naïve, non-severe |
| 15 | 35% | Yes | Naïve, severe |
| 16 | 8% | No | Naïve, non-severe |
| 17 | 2.5% | No | Pre-exposed, non-severe |
| 18 | 7% | No | Pre-exposed, non-severe |
| 19 | 0.8% | No clinical data | Pre-exposed, non-severe |
| 20 | 40% | No | Naïve, severe |
| 21 | 7% | No | Naïve, non-severe |
| 22 | 3% | No | Naïve, non-severe |
| 23 | 12% | Yes | Naïve, severe |
| 24 | 3% | No | Pre-exposed, non-severe |
| 25 | 48% | Yes | Naïve, severe |
| 26 | 7% | Yes* | Naïve, severe |
| 27 | 0.2% | No | Pre-exposed, non-severe |
| 28 | 3.5% | No | Pre-exposed, non-severe |
| 29 | 8% | No | Pre-exposed, non-severe |
| 30 | 11% | Yes | Naïve, severe |
| 31 | 1% | No clinical data | Pre-exposed, non-severe |
| 32 | 2% | No | Pre-exposed, non-severe |
| 33 | 3.5% | No | Pre-exposed, non-severe |

* circulating schizonts upon hospitalization
